# Kinetic and functional analysis of abundant microRNAs in extracellular vesicles from normal and stressed cultures of Chinese Hamster Ovary (CHO) cells

**DOI:** 10.1101/2023.06.21.545902

**Authors:** Jessica Belliveau, Will Thompson, Eleftherios Terry Papoutsakis

## Abstract

Chinese hamster ovary (CHO) cells release and exchange large quantities of extracellular vesicles (EVs). EVs are highly enriched in microRNAs (miRs, or miRNAs), which are responsible for most of their biological effects. We have recently shown that the miR content of CHO EVs varies significantly under culture stress conditions. Here, we provide a novel stoichiometric (“per-EV”) quantification of miR and protein levels in large CHO EVs produced under ammonia, lactate, osmotic, and age-related stress. Each stress resulted in distinct EV miR levels, with selective miR loading by parent cells. Our data provide a proof of concept for the use of CHO EV cargo as a diagnostic tool for identifying culture stress. We also tested the impact of three select miRs (let-7a, miR-21, and miR-92a) on CHO cell growth and viability. Let-7a—abundant in CHO EVs from stressed cultures—reduced CHO cell viability, while miR-92a—abundant in CHO EVs from unstressed cultures—promoted cell survival. Overexpression of miR-21 had a slight detrimental impact on CHO cell growth and viability during late exponential-phase culture, an unexpected result based on the reported anti-apoptotic role of miR-21 in other mammalian cell lines. These findings provide novel relationships between CHO EV cargo and cell phenotype, suggesting that CHO EVs may exert both pro- and anti-apoptotic effects on target cells, depending on the conditions under which they were produced.

**Graphical Abstract:** 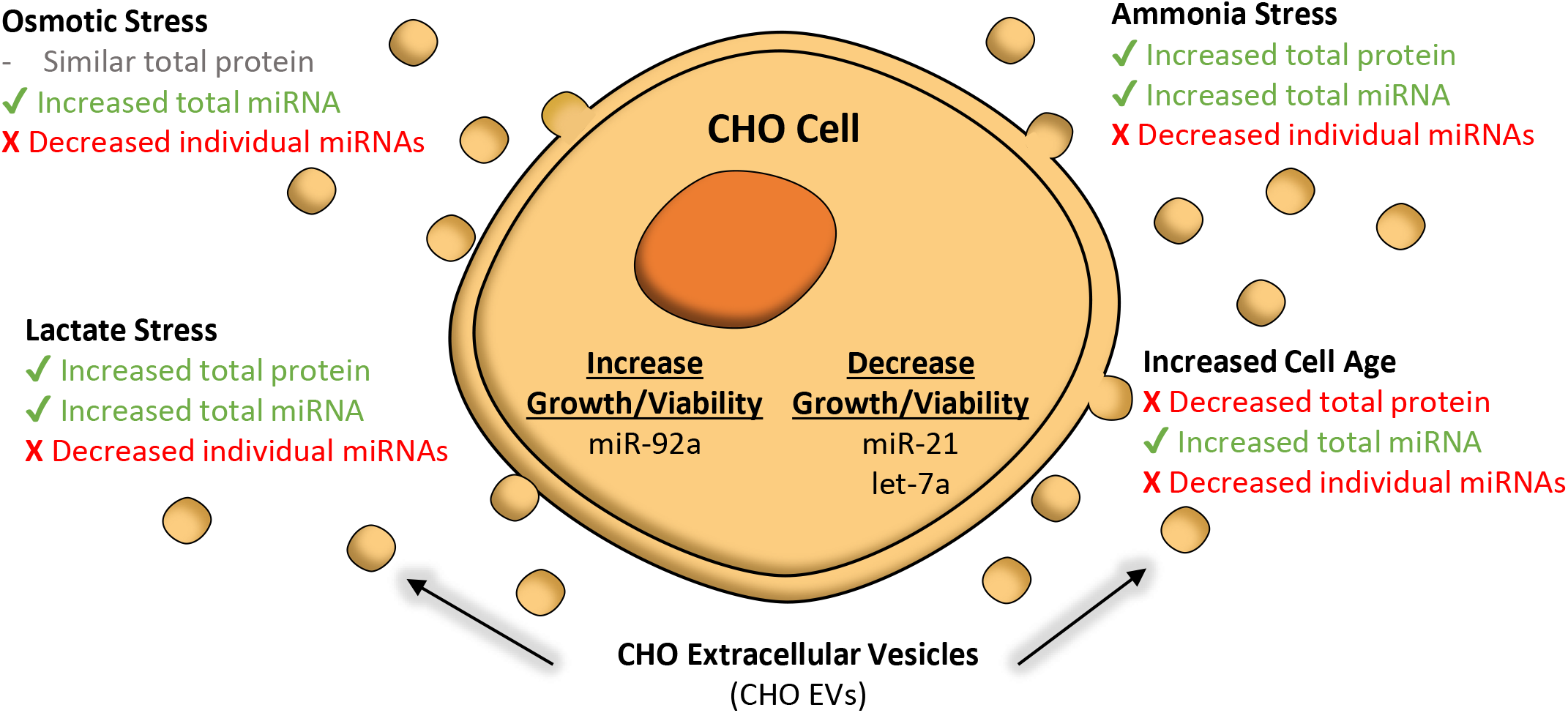

## 1. INTRODUCTION

Chinese hamster ovary (CHO) cells have long served as mainstays of industrial protein production. Recently, we have found that CHO cells in culture participate in a massive and continuous exchange of extracellular vesicles (EVs) (Belliveau & Papoutsakis, 2022), thus impacting culture performance by a hitherto unknown and unexplored mechanism. EVs are submicron-sized particles enclosed by a lipid bilayer which originate from either multivesicular bodies or the plasma membrane. EVs from multivesicular bodies (known as “exosomes”) are generally smaller in size (50-150 nm), while EVs released from the plasma membrane (known as microparticles or microvesicles) tend to be larger (100-1,000 nm). Released by every cell type, EVs serve as linchpins of intercellular communication, mediating phenotypes of target cells through the delivery of nucleic acid and/or protein cargo (Kao & Papoutsakis, 2019; Raposo & Stoorvogel, 2013; van Niel, D’Angelo, & Raposo, 2018).

In particular, microRNAs (miRs) have been identified as key molecules responsible for EV biological activities (Dellar, Hill, Melling, Carter, & Baena-Lopez, 2022; O’Brien, Breyne, Ughetto, Laurent, & Breakefield, 2020). Most commonly, miRs influence cellular behavior by binding to complementary sequences on longer mRNAs and subsequently inhibiting protein translation or degrading the mRNA (Dellar et al., 2022; O’Brien et al., 2020). Most miRs have multiple mRNA targets and work together to regulate cellular processes and phenotypes; even small changes in miR expression, such as 1.5-fold change, can affect cellular phenotypes (Mestdagh et al., 2009). EVs are loaded with individual miRs and other cargo by a variety of chaperone proteins (Anand, Samuel, Kumar, & Mathivanan, 2019; Corrado, Barreca, Zichittella, Alessandro, & Conigliaro, 2021; Fabbiano et al., 2020; Leidal & Debnath, 2020).

CHO EVs have only recently garnered significant attention. In particular, we have found that CHO EVs exchange large quantities of RNA cargo between their parent cells (Belliveau & Papoutsakis, 2022, 2023). The nature of that cargo, which has been investigated by several groups (Belliveau & Papoutsakis, 2023; Busch et al., 2022; Keysberg et al., 2021), is now known to vary significantly as a function of metabolite (ammonia and osmotic) stress (Belliveau & Papoutsakis, 2023). Preliminary proteomic analysis has also been performed on CHO EVs, with large and diverse protein populations observed in both large and small CHO EVs from various phases of culture (Keysberg et al., 2021; Kumar et al., 2016). Still, nothing is known about the specific function of CHO EV cargo in culture. In two instances, CHO EVs have been found to promote desirable phenotypes in target cells, with smaller EVs (i.e., “exosomes”) promoting CHO cell growth (Takagi, Jimbo, Oda, Goto, & Fujiwara, 2021) and protecting CHO cells against oxidative stress (Han & Rhee, 2018). However, the EV cargo mediating these phenotypes remains unknown.

From this vantage point, this study provides novel insights into the functions of CHO EV miR cargo in CHO cells, hinting at possible mechanisms for CHO EV bioactivity observed in other studies. Specifically, we provide a unique (for CHO EV research) stoichiometric accounting of miR and protein cargo levels in EVs produced under various stress conditions. We also examine cellular phenotypes mediated by the three highly-abundant miRs: let-7a, miR-21, and miR-92a. The focus of this work is the larger EVs originating largely from the plasma membrane (i.e., CHO “microparticles,” or “CHO MPs”), which have been the focus of fewer studies, but which are easier to isolate—a boon for their potential as a diagnostic tool—and capable of carrying larger cargo loads, thus suitable for a broader range of synthetic applications using DNA, RNA, protein, nucleoprotein (such as CRISPR systems) and other organic molecules as payload. Our previous publications (Belliveau & Papoutsakis, 2022, 2023) have provided detailed CHO MP characterization that meets or exceeds the MISEV2018 guidelines from the International Society for Extracellular Vesicles (Thery et al., 2018). Here, we use the same isolation methods as before, expanding our CHO MP characterization by focusing primarily on various miR cargoes and their potential functions in CHO cell culture.

## 2. MATERIALS AND METHODS

### 2.1 Chemicals and reagents

Except where otherwise noted, all chemicals and reagents were obtained from Sigma-Aldrich (St. Louis, MO, USA) or Thermo Fisher Scientific (Waltham, MA, USA).

### 2.2 CHO cell culture

CHO cells were obtained and cultured as described previously (Belliveau & Papoutsakis, 2022, 2023). Briefly, a CHO-K1 cell line expressing the VRC01 antibody was cultured in HyClone ActiPro media (Cytiva, Marlborough, MA, USA) supplemented with 6 mM L-glutamine. Culture occurred in either 125 mL shake flasks (20 mL culture volume) at 120 rpm or 50 mL culture tubes (15 mL culture volume) at 250 rpm. A seeding density of 0.4 x 10^6^ cells/mL was employed in each case, and cells were incubated at 37°C in 20% O_2_ and 5% CO_2_.

Ammonia stress was applied by adding 9 mM ammonium chloride (NH_4_Cl) in phosphate-buffered saline (PBS) (pH 7.4) to day 0 of the culture. Lactate stress was applied by adding 10 mM sodium lactate (NaC_3_H_5_O_3_) in PBS (pH 7.4) to day 2 of the culture (to coincide with the natural spike in culture lactate concentration). Osmotic effects of the lactate stress were controlled for via an “osmotic control” treatment, wherein 10 mM of sodium chloride (NaCl) in PBS (pH 7.4) was added to day 2 of the culture, such that the culture osmolarity resulting from the added NaCl was 20 mOsm/L. Higher osmotic stress was applied by adding 60 mM sodium chloride (NaCl) in PBS (pH 7.4) to day 0 of the culture, such that the culture osmolarity resulting from the added NaCl was 120 mOsm/L.

### 2.3 Isolation and quantification of extracellular vesicles

CHO EVs were isolated as described (Belliveau & Papoutsakis, 2022, 2023). As noted, only large CHO EVs (i.e., “CHO MPs”) were collected and analyzed for this study. Briefly, CHO cells were pelleted via centrifugation at 180 x g for 4 min. From the resulting supernatant, apoptotic bodies and large cellular debris were pelleted via centrifugation at 2,000 x g for 10 min. Finally, from this new supernatant, CHO MPs were pelleted via ultracentrifugation at 28,000 x g and 4°C for 30 min. Ultracentrifugation employed an Optima LE-80K Ultracentrifuge with an SW-28 rotor (Beckman Coulter, Brea, CA, USA) for initial separation and an Optima Max Ultracentrifuge with a TLA-55 rotor (Beckman Coulter, Brea, CA, USA) for sample enrichment and washing. Isolated CHO MPs were resuspended in culture media and used immediately or stored at 4°C overnight.

CHO MPs were counted using either a BD FACSAriaII flow cytometer with FACSDiva software (BD Biosciences, Franklin Lakes, NJ, USA) or a CytoFLEX S flow cytometer with CytExpert software (Beckman Coulter, Brea, CA, USA). Here, CHO MPs were defined as all particles between 0.2 micron (the smallest reliable size that can be counted using our flow cytometry instruments) and 1 micron in size; size gates were constructed using fluorescent calibration beads from Spherotech (Lake Forest, IL, USA). For experiments using the BD FACSAriaII flow cytometer, a known quantity of ∼5 micron AccuCount Fluorescent Beads (Spherotech, Lake Forest, IL, USA) was used for CHO MP counting.

### 2.4 Extraction of total RNA

Extraction of total RNA from CHO cells and MPs was done as described (Belliveau & Papoutsakis, 2023). Briefly, total RNA was extracted from pelleted cells or MPs using a miRNeasy Mini Kit (QIAGEN, Hilden, Germany) per manufacturer’s instructions. In some cases, synthetic cel-miR-39-3p (Thermo Fisher Scientific, Waltham, MA, USA) was added to samples during the cell/MP lysis step as a spike-in control. Extracted RNA samples were flash frozen in liquid N_2_ and stored at -80°C until further use.

### 2.6 Quantification of total miR

Total miR levels in extracted RNA samples were quantified using a Qubit miR Assay Kit and Qubit 3.0 Fluorimeter (Thermo Fisher Scientific, Waltham, MA, USA); the manufacturer’s protocols were employed.

### 2.5 Quantification of individual miRs via RT-PCR

Individual miR levels were quantified via RT-PCR as described (Kao, Jiang, Thompson, & Papoutsakis, 2022). Briefly, reverse transcription employed the ThermoFisher TaqMan MicroRNA Reverse Transcription Kit with miR-specific primers and probes (Thermo Fisher Scientific, Waltham, MA, USA), and proceeded according to the manufacturer’s protocol. PCR employed the TaqMan Universal PCR Master Mix II with miR-specific Small RNA Assays (Thermo Fisher Scientific, Waltham, MA, USA), and proceeded according to the manufacturer’s protocol. For PCR experiments, three technical replicates were performed for each biological replicate. Reverse transcription and PCR used a CFX96 Optical Reaction Module (Bio-Rad, Hercules, CA, USA), with miR levels quantified via the 2^-ΔΔCT^ method (Livak & Schmittgen, 2001).

### 2.6 Quantification of total protein

Total protein in CHO MP samples was quantified using a Bradford-based, colorimetric assay kit. CHO MPs were washed in PBS and resuspended in RIPA buffer (Millipore Sigma, Burlington, MA, USA) supplemented with protease inhibitor (200:1 buffer-to-inhibitor ratio). Sample volumes ranged from 50 to 300 μL, depending on CHO MP concentration. Samples were agitated at 4°C for 30 min. and thereafter centrifuged at 16,000 x g and 4°C for 25 min. Supernatants containing the total protein were diluted with water (18:2 water-to-lysate ratio). Subsequent protein quantification employed the Bio-Rad protein assay kit (Bio-Rad, Hercules, CA, USA). A 100 μL solution of Reagent A and Reagent S (5:1 ratio) was added to each sample; samples were agitated briefly and incubated at room temperature for 15 min. Absorbance of each sample at 750 nm was recorded for three technical replicates of each biological replicate. A calibration curve was constructed via serial dilution of a BSA standard.

### 2.7 Plasmid preparation

*E. coli* transformed with plasmids, detailed below, containing the primary or a fragment of the primary miR sequences were obtained from Addgene as stab cultures, expanded on LB agar plates with 100 μg/mL carbenicillin, and single colonies were selected to expand in liquid LB cultures with 100 μg/mL carbenicillin overnight. Plasmid purification of the overnight cultures was done with QIAGEN Miniprep Kit (QIAGEN, Hilden, Germany) and quantified with the dsDNA HS Qubit Kit (Invitrogen, Waltham, MA, USA).

### 2.8 Overexpression of miRs

CHO cells were electroporated with 2 μg of plasmids containing the primary or a fragment of the primary microRNA sequences of three miRs that were in high abundance in standard or stressed cultures (miR-21 (Addgene #21114), miR-92a (Addgene #46672), let-7a (Addgene #51377) using the Nucleofector V kit (Lonza, Basel, Switzerland) according to manufacturer’s protocols. Cells were cultured in 2 mL cultures in 12-well plates at 37°C and 120 RPM. Growth media was changed one day after electroporation to remove excess electroporation buffer and 0.2 mg/mL geneticin was added to apply selection pressure to overexpress the plasmids. Geneticin was added every other day to maintain selection pressure. Cell growth was measured every day and viability was measured every other day.

### 2.9 Viability assay

Viability was measured every other day using an Annexin V (BioLegend, San Diego, CA, USA) and 7AAD (BioLegend, San Diego, CA, USA) flow cytometry assay. Briefly, cells were washed with cold PBS twice, then resuspended in Annexin V Binding Buffer (BioLegend, San Diego, CA, USA) and incubated at room temperature in the dark with FITC-Annexin V (1:20 antibody to binding buffer by volume) and 7AAD (1:20 stain to binding buffer by volume) for 15 minutes. Cells were then analyzed using flow cytometry (CytoFLEX S flow cytometer using CytExpert Software, Beckman Coulter, Brea, CA, USA).

### 2.10 Statistical analysis

Except where otherwise noted, each data point represents the mean of ≥3 biological replicates, with error bars indicating ± one standard error of the mean. Unpaired Student’s t-tests were performed on all data, with * indicating p<0.05, ** indicating p<0.01, and *** indicating p<0.001.

## 3. RESULTS

### 3.1 Total miR and protein quantities in CHO EVs vary with culture age and stress

Total miR and protein levels in CHO EVs could serve as valuable metrics for CHO culture diagnostics but also for cell-specific miR manufacturing/production. For example, in vitro or in vivo dosages of EVs for biotechnological or therapeutic applications are based on total mass or total proteins, two quantities that are affected by the specific per EV mass or protein. How those might vary with culture parameters has not been systematically reported. Although our prior work has identified individual miR cargo in CHO EVs produced under stress (Belliveau & Papoutsakis, 2023), nothing is known about age/stress-induced variation in total miR or protein levels in CHO EVs. In this study, total miR and protein levels were quantified using Qubit fluorimetry and a Bradford protein assay, respectively. Total miR levels in CHO MPs increased steadily as parent cultures aged, with significant increases in said miR levels occurring from day 3 to day 6 and from day 6 to day 9 (Fig 1a). For CHO MP protein levels, however, the trend was reversed: levels were highest on day 1 and diminished thereafter, with protein levels dropping significantly from day 3 to day 6 and from day 6 to day 9 (Fig 1b). Notably, total protein weight is much higher than total miR weight at any given point in time. These results were unexpected and surprising in that one would assume that these two EV specific properties are largely constant. The opposite trends (total miR vs total protein) are also unexpected and the mechanism that underlies that is for now unknown.

**Figure 1.**
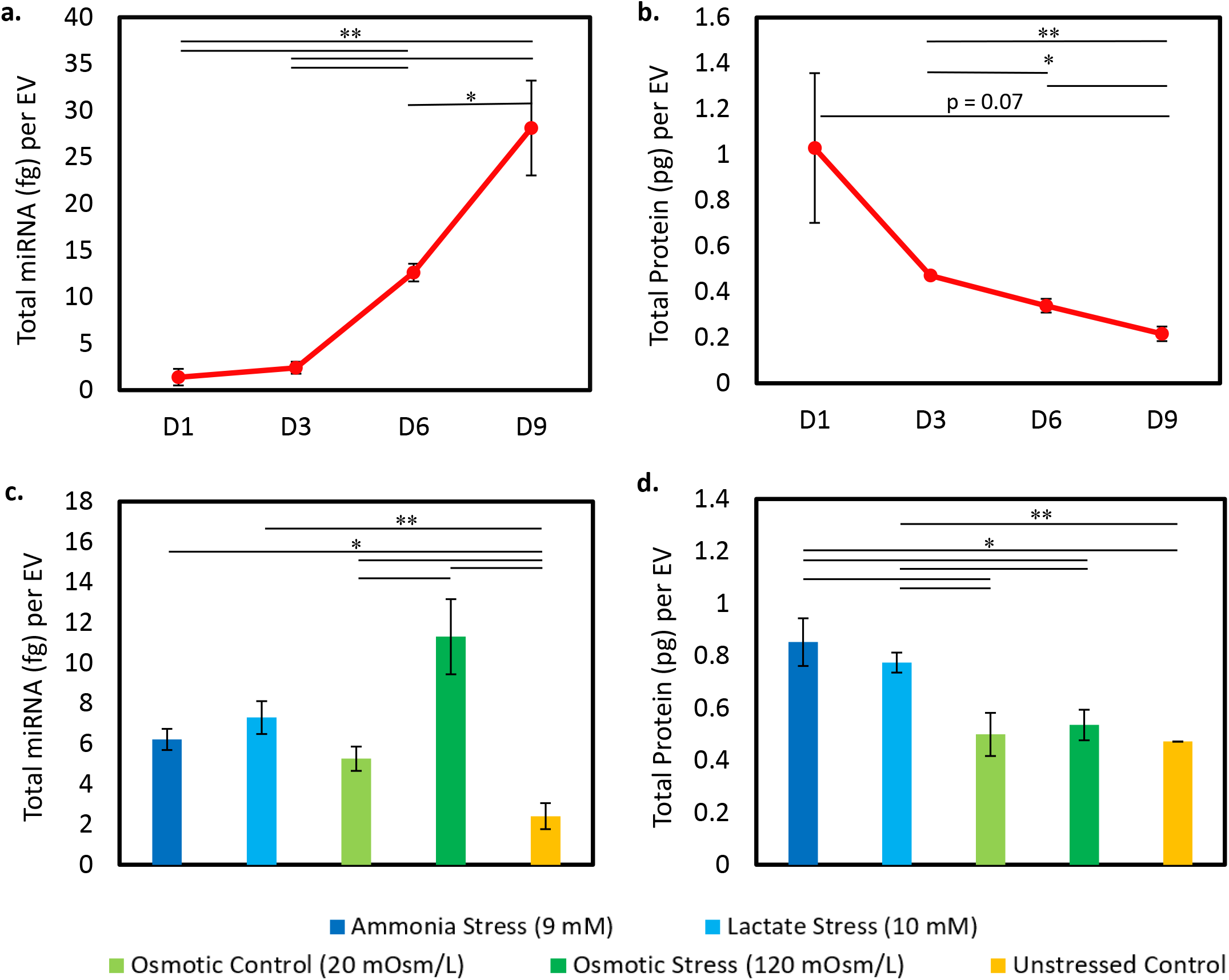
Total miR and protein levels in CHO MPs. Total **(a)** miR and **(b)** protein levels in CHO MPs were measured at different points in culture and normalized to CHO MP quantity. Total **(c)** miR and **(d)** protein levels in CHO MPs produced under osmotic, ammonia, and lactate stress were measured and normalized to CHO MP quantity. Error bars represent the standard error of the mean (SEM) of 3 replicates. Significance was determined with unpaired Student’s t-test, * for p<0.05, ** for p<0.01.

Total miR levels in CHO MPs were also significantly higher (relative to controls) in day 3 cultures treated with ammonia, lactate, or osmotic (20 and 120 mOsm/L) stress. 120 mOsm/L osmotic stress was also significantly more effective than 20 mOsm/L osmotic stress in boosting total CHO MP miR levels (Fig 2c). However, no level of osmotic stress had an impact on total CHO MP protein levels; only ammonia and lactate stress promoted significant increases in total protein levels (relative to the control) (Fig 2d). In sum, these results suggest that miR and protein loading in CHO EVs is a dynamic process that varies significantly with culture age and stress. While total miR or protein levels are inappropriate as methods for CHO EV counting, they offer promise as metrics for the rapid and non-invasive monitoring of culture stress conditions.

**Figure 2.**
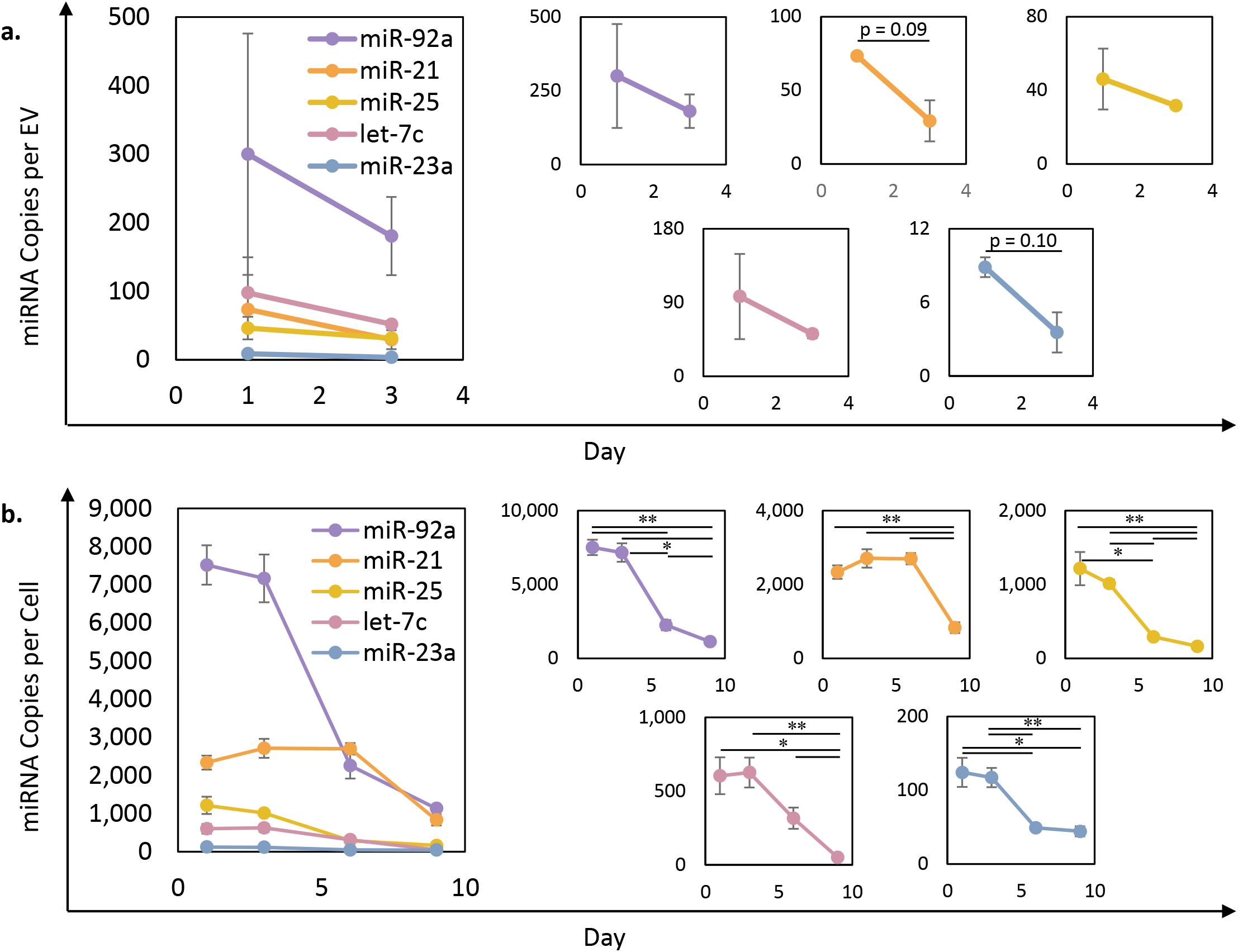
Individual miR levels in CHO MPs and CHO cells at various timepoints in culture. The levels of five abundant miRs were individually measured in **(a)** CHO MPs and **(b)** CHO cells at different points in culture via RT-PCR. Error bars represent the standard error of the mean (SEM) of 3 replicates. Significance was determined with unpaired Student’s t-test, * for p<0.05, ** for p<0.01.

### 3.2 Levels of highly abundant miRs in CHO cells decrease with culture age

Given the relationship between culture age and CHO EV miR cargo loading, optimal harvest times for CHO EVs with high levels of specific individual miRs must be identified, such that per-EV miR levels are maximized. In this study, RT-PCR was employed to quantify levels of five highly abundant miRs (as determined by RNA sequencing of CHO MPs from standard culture) (Belliveau & Papoutsakis, 2023). Levels of cgr-miR-92a-3p, cgr-miR-23a-3p, cgr-miR-21-5p, cgr-miR-25-3p, and mmu-let-7c-5p were assessed in both CHO MPs and CHO cells on days 1, 3, 6 and 9 of culture. Highly sensitive TaqMan assays combined with spike-in control cel-miR-39-3p enabled calculation of miR copy number per EV via the 2^-ΔΔCT^ method (Livak & Schmittgen, 2001), which is an increasingly popular method for EV miR cargo quantification (Kondratov et al., 2020; Perge et al., 2017). Cellular levels of all five miRs decreased significantly as cultures aged, with the most significant drops observed between day 3 and day 9 (Fig 2b). Similar decreases were observed between CHO MPs derived from day 1 and day 3 cultures, though these drops were not significant (Fig 2a). Individual miR levels in CHO MP samples from days 6 and 9 were not detectable via RT-PCR, possibly the result of a co-isolated product inhibiting reverse transcription and/or amplification. Nevertheless, trends in cellular miR levels suggest that CHO MP miR levels may also vary significantly with culture age. Optimization of harvest times for CHO EVs possessing high or low levels of various specific miRs must therefore be a priority.

### 3.3 Highly abundant individual miRs are more highly enriched in CHO MPs from unstressed cultures, a phenomenon not observed in the parent cells

Levels of the individual miRs noted above were also measured in CHO MPs (Fig 3a) and CHO cells (Fig 3b) produced/grown under osmotic, ammonia, and lactate stress. Samples were taken from day 3 culture, with samples from unstressed cultures serving as controls. Since lactate stress did not inhibit cell growth, a 20 mOsm/L NaCl control was used to assess whether the impact of lactate on EV miR cargo was due to its osmotic effects. The individual miR levels in CHO MPs from unstressed cultures were, without exception, higher than the levels observed in CHO MPs from stressed cultures, regardless of the type or magnitude of stress, and this trend was statistically significant for miR-25 and let-7c. Notably, despite these relatively large differences between individual miR quantities in stressed and unstressed CHO MPs, miR quantities rarely varied significantly between stressed unstressed parent cells (Fig 3b), suggesting the impact of stress on CHO MP miR levels is mediated not by cellular miR concentrations, but rather by changes in the behavior of protein chaperones responsible for cargo loading (for the five miRs and four stress conditions tested, at least). On the other hand, cellular miR levels under ammonia/lactate stress often were significantly higher than corresponding levels under osmotic stress (Fig 3b), suggesting the various stresses have highly distinct impacts on cellular miR production machinery.

**Figure 3.**
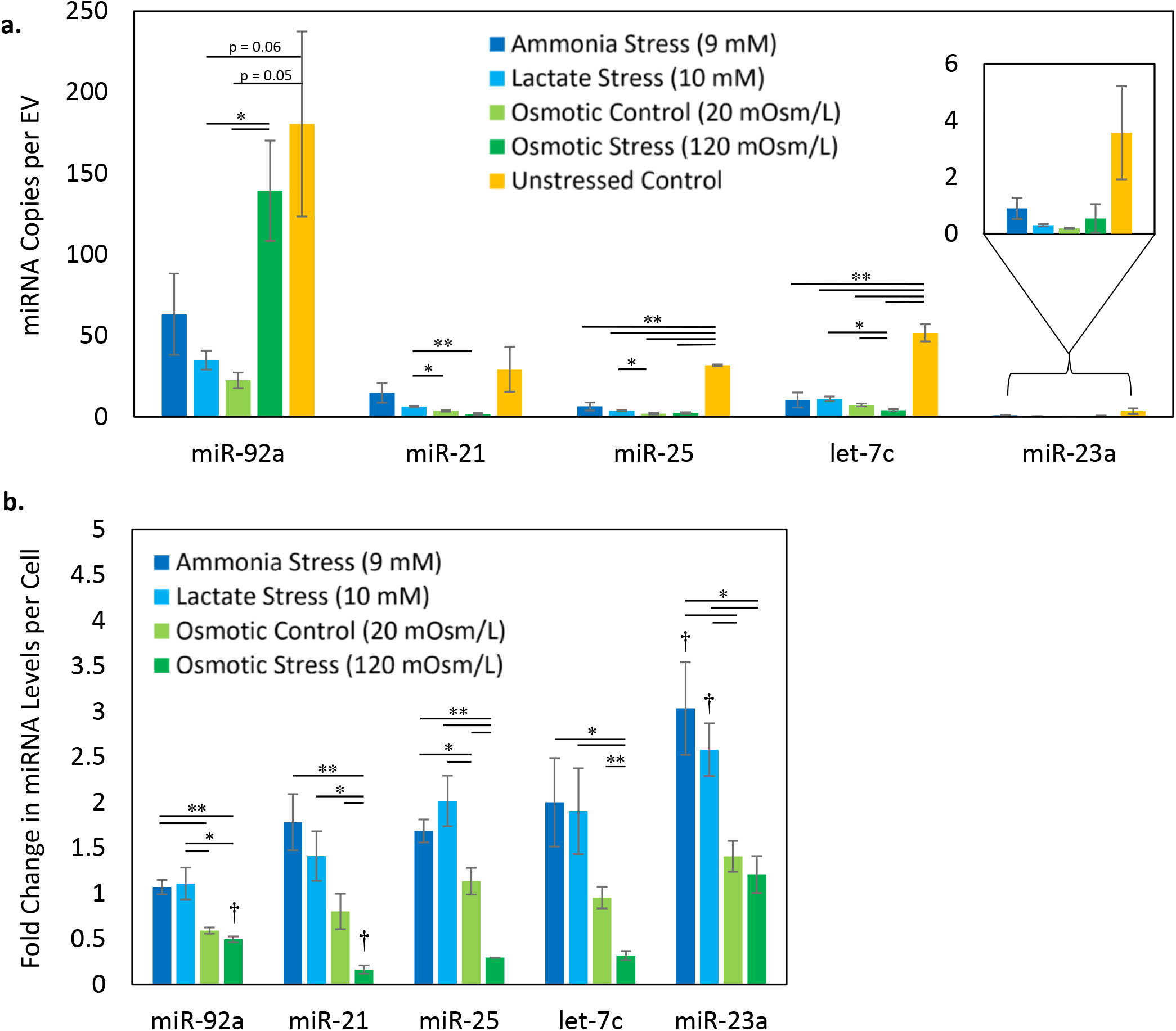
Individual miR levels in CHO MPs and CHO cells from stressed cultures. The levels of five abundant miRs were individually measured in **(a)** CHO MPs and **(b)** CHO cells from stress-treated day 3 cultures via RT-PCR. The bars in (b) represent fold change relative to unstressed control cell miR levels. Error bars represent the standard error of the mean (SEM) of 3-4 replicates. Significance was determined with unpaired Student’s t-test. For significance relative to another stressed sample, * for p<0.05, ** for p<0.01; for significance relative to the unstressed control, † for p<0.05.

Previous investigations of EVs from other cell types have found abundant miR species to exist in EVs at anywhere from 1 copy per 10–100 EVs (Chevillet et al., 2014; Wei et al., 2017) to 10–100 copies per 1 EV (Kondratov et al., 2020; Stevanato, Thanabalasundaram, Vysokov, & Sinden, 2016). The individual miR copy numbers (per EV) identified in this study fit quite reasonably within these ranges, with especially notable expression (>100 copies) of miR-92a per EV observed in two samples. Traditional flow cytometry is limited in its ability to count particles of less than 200 nm, while nanoparticle tracking analysis (NTA) accounts for smaller particles but is more apt to count non-EV proteins and aggregates (George et al., 2021). For this reason, EV concentrations are 1-2 orders of magnitude higher when counts are taken via NTA (versus flow cytometry) (George et al., 2021). Given our focus on larger MPs, which are enriched via ultracentrifugation at 28,000 x g, this study counts EVs using flow cytometry.

The RT-PCR data in this section also provide support for a selective CHO MP loading process, both during normal growth and under stress conditions. That is, the miR contents of the CHO MPs are not a proportional reflection of intracellular miR availability, but instead reflect an active, intentional loading process on the part of the parent cell and its protein chaperones. Assuming spherical CHO cells of 15 microns (diameter) and spherical CHO MPs of 250 nm, CHO MPs from unstressed (control) day 3 culture contain volumetric concentrations of miR-92a, miR-21, miR-25, let-7c, and miR-23a that are 5,400, 2,300, 6,800, 18,000, and 6,600 times greater, respectively, than those in their parent cells. This exceptionally high enrichment of miR would logically be interpreted to mean that the parent cells may achieve specific biological goals by this enrichment, and since these large EVs are largely budding off the cytoplasmic membrane that the cells concentrate specific miRs near the cytoplasmic membrane to enable this highly concentrated miR loading.

### 3.4 Transient overexpression of individual miRs impacts CHO cell growth and viability

Little is currently known regarding the functions of CHO EV miR cargo in target CHO cells. Therefore, we wanted to examine the growth, viability, and apoptosis of CHO cells overexpressing either let-7a, which is highly expressed in CHO MPs from stressed cultures, or miR-21 or miR-92a, both of which are highly expressed in CHO MPs from unstressed, exponential-phase cultures (Belliveau & Papoutsakis, 2023). Let-7a has been reported in the literature to have a negative impact on cell growth and viability in other cell types (Cho, Song, Oh, & Lee, 2015; Tsang & Kwok, 2008; Zhao et al., 2018). On the other hand, miR-21 and miR-92a have been reported to improve cell growth and viability (Feng & Tsao, 2016; Krichevsky & Gabriely, 2009; Liu, Wang, Yang, Xiao, & Chen, 2014; Mogilyansky & Rigoutsos, 2013; Shigoka et al., 2010; Thabet et al., 2020; Xu et al., 2019; Yang et al., 2019; Zhou et al., 2015). In CHO cells, miR-92a has also been shown to boost protein production by raising intracellular cholesterol levels (Loh, Yang, & Lam, 2017).

In this study, cultures were transiently transfected with plasmids encoding miRs that were previously identified to be in high abundance during exponential phase in stressed or standard cultures (let-7a, miR-21, miR-92a) to observe a relationship between these specific miRs and cell growth and viability. We hypothesized overexpressing highly abundant miRs from unstressed cultures identified in exponential phase, miR-21 and miR-92a, throughout the culture lifespan would promote cell growth and viability, particularly in stationary phase. Conversely, we hypothesized that overexpression of highly abundant miRs from stressed cultures, let-7a, would decrease cell growth and viability. Cultures were compared to a control culture that was transiently transfected with a plasmid encoding a far-red fluorescent protein (pLifeAct-miRFP703) to control for the effects of electroporation, plasmid burden, and selection pressure. The burden of expressing a protein (1 kb) is greater than that of expressing pri-miR sequences (80-150 bp) and, in addition, the expressed protein is a fluorescent reporter that does not have a functional role in the cell. Therefore, the differences in the plasmid effects on cell growth and viability between this control and miR-overexpressing conditions are likely conservative. Another type of control that could have been used is a plasmid with a “random sequence” miR, or an “empty” plasmid (i.e., a plasmid without DNA in the locus of the miR). While widely tested for human miRs, we are concerned that one can never ascertain the a “random sequence” miR has not biological effects in CHO cells for lack of extensive testing. The “empty” option would be preferrable, but in the end we opted for the more conservative pLifeAct-miRFP703 control.

In cultures that were transiently transfected with the let-7a plasmid (Figure 4a), the cell concentrations at days 3, 6, 7, 8, and 9 were significantly lower compared to the control culture, indicating let-7a has a negative impact on cell growth. The cell density of cultures transfected with the plasmid encoding miR-21 was only significantly different from the control culture at days 7 and 8. For cultures with the miR-92a plasmid, the cell density was significantly different from the control on day 2; otherwise, there was no difference in the cell growth curve. The viability of these cultures was measured on days 4, 6, and 8 with an annexin V/7AAD flow cytometry assay (Figure 4b). Conditions in red indicate reduced viability or increased apoptosis and conditions in green indicate increased viability or decreased apoptosis compared to the control culture at the same timepoints. On days 4 and 6, cultures transfected with let-7a had significantly fewer viable cells. The let-7a condition had significantly more early apoptotic and late apoptotic cells compared to the control culture, in agreement with reports of let-7a inducing apoptosis (Cho et al., 2015; Tsang & Kwok, 2008).

**Figure 4.**
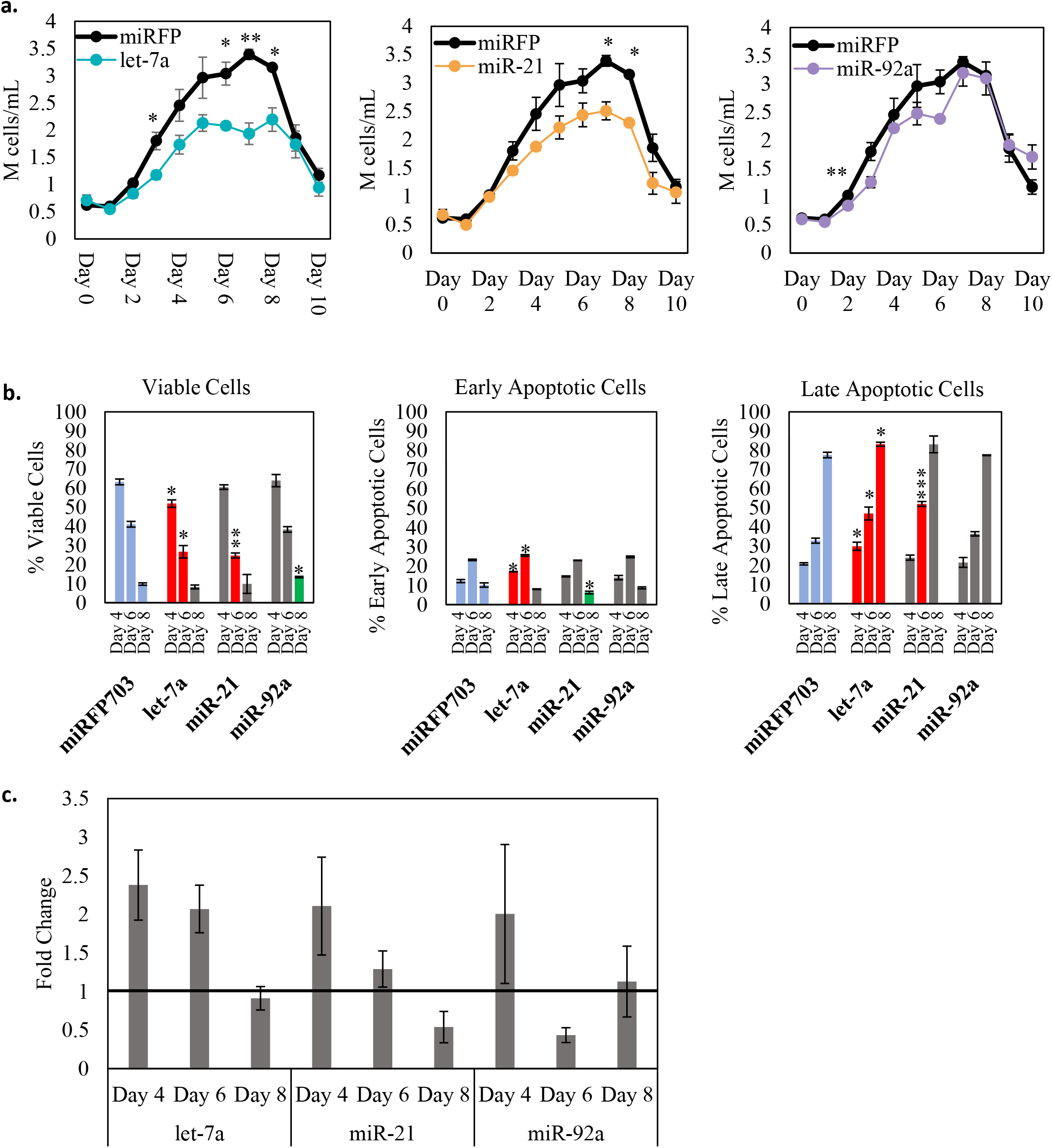
Transient expression of let-7a, miR-21, and miR-92a. **(a)** Cell growth curve of cells transiently expressing let-7a, miR-21, or miR-92a compared to the control (transient transfection of pLifeAct-miRFP703). **(b)** Annexin V/7AAD flow cytometry viability assay of cultures transiently transfected with the control plasmid (pLifeAct-miRFP703), let-7a plasmid, miR-21 plasmid, or miR-92a plasmid. Conditions where viability was significantly lower than the associated control are red. Conditions where viability was significantly greater than the associated control are green. **(c)** Fold change in cultures transiently expressing let-7a, miR-21, and miR-92a compared to the endogenous expression in the control culture (transient transfection of pLifeAct-miRFP703). Error bars represent the standard error of the mean (SEM) of 3-4 replicates. Significance relative to control was determined with unpaired Student’s t-test, * for p<0.05, ** for p<0.01, *** for p<0.001.

Cultures with the plasmid expressing miR-21 demonstrated decreases in the populations of viable and early apoptotic cells and an increase in the population of late apoptotic cells on day 6 of culture. miR-21 is highly expressed during exponential phase in normal batch cultures and is reported in other mammalian cell lines to have an anti-apoptotic role and support cell growth (Feng & Tsao, 2016; Krichevsky & Gabriely, 2009); therefore, we initially hypothesized that overexpression of miR-21 through stationary phase would support both cell growth and viability. However, in cultures overexpressing miR-21, a decrease in cell density and a decrease in cell viability was observed during late stationary phase. Our study therefore suggests that the impacts of overexpressing miR-21 by up to twofold on CHO cell growth and viability are negligible in early culture, and may be negative in mid to late culture. Cultures with the plasmid expressing miR-92a demonstrated an increase in the population of viable cells on day 8 of culture compared to the control culture at the same timepoint. In the literature, miR-92a is reported to increase cell proliferation (Zhou et al., 2015), and was observed in our previous experiments to be highly expressed early in culture (Figure 2). Here, overexpression of miR-92a resulted in improved viability in late culture.

The fold change in miR levels, measured via RT-qPCR, was expressed relative to endogenous expression in a control culture with transient transfection of apLifeAct-miRFP703 plasmid (Figure 4c). Approximately a 2-fold difference in expression of let-7a, miR-21, and miR-92a was observed on day 4 of culture, which is well within the range generally required for phenotypic changes (Mestdagh et al., 2009).

Taken together, these results suggest novel pro- and anti-apoptotic roles for highly-expressed CHO MP miR cargo in target CHO cells. The let-7a data suggest a potential pro-apoptotic function for CHO MPs produced under ammonia/osmotic stress conditions (which upregulate let-7a cargo levels). The miR-92a data suggest a potential anti-apoptotic function for CHO MPs produced in unstressed, exponential-phase cultures (which upregulate miR-92a cargo levels). Indeed, it is possible that miR-92a action plays a role in the protective effects of CHO EVs reported in other studies (Han & Rhee, 2018; Takagi et al., 2021). The miR-21 data in this study were unexpected, because, as noted previously, miR-21 has repeatedly been found to promote growth and viability of other cell types.

## 4. DISCUSSION

In this study, we measured total miR and protein levels on a stoichiometric (“per-EV”) basis, and then used RT-PCR to quantify the presence of total miR and protein cargo, as well as key individual miRs. Total miR levels were significantly higher in CHO MPs from stressed cultures (Fig 1c), even as individual miR levels were often significantly lower (Fig 3a). This finding suggests that the bulk of the total miR content in CHO MPs from stressed cultures is *not* composed of the five miRs we tested individually (i.e., the five most abundant miRs in CHO MPs from control culture). Indeed, RNA sequencing partly confirms this hypothesis, as identities of the most abundant miRs in CHO MPs differ when parent cells are exposed to ammonia and osmotic stress (Belliveau & Papoutsakis, 2023). We would expect a similar finding following the application of lactate stress. Similarly, we expect that RNA sequencing of CHO MPs from different growth phases would detect distinct miR profiles (relative to CHO MPs from day 3 culture) (Keysberg et al., 2021), since CHO MPs under these conditions carry more total miR (Fig 1a) despite an apparent downward trend in individual miR levels during early culture (Fig 2a). The stoichiometric analysis of per-EV miR and protein levels presented in this study is unique in CHO EV research, which has so far been limited to descriptions of relative RNA levels informed by RNA sequencing data.

Notably, this study represents the first analysis of the impact of lactate stress on CHO EV cargo. Although the lactate concentrations employed in this study did not have a significant impact of cell growth or viability, miR and protein levels in EVs were certainly affected. Lactate appears similar to ammonia in terms of its effect on miR and protein cargo, and its impact extends beyond its osmotic effects, which were controlled for by a 20 mOsm/L NaCl treatment (i.e., the “osmotic control”).

This study also explored the role of three specific miRs (let-7a, miR-21, and miR-92a) on cell growth, viability, and apoptosis in CHO cultures across growth phases. In transiently overexpressing let-7a, there was an observed decrease in cell growth and viability. There was no impact to cell growth in cultures transfected with the miR-92a plasmid and there was an observed increase in viability of these cultures on days 2 and 8. Cultures transfected with the miR-21 plasmid did not demonstrate improved cell growth or viability (such improvement was expected, based on literature reports), with decreases in viability only on day 6 of culture. This result was somewhat surprising, and miR-21 has been associated with improved growth and viability in other cell types (Feng & Tsao, 2016; Krichevsky & Gabriely, 2009; Liu et al., 2014; Thabet et al., 2020; Yang et al., 2019).

These results suggest, for the first time, potential functions for the common miR cargoes carried by CHO EVs from stressed and unstressed cultures. Namely, let-7b (common in CHO EVs from stressed cultures) may induce CHO cell apoptosis, while miR-92a (common in CHO MPs from unstressed, exponential-phase cultures) may promote CHO cell survival. A more robust study stably overexpressing these miRs in CHO cells will be required to fully understand the role these miRs have on industrially-relevant parameters such as cell growth, viability, apoptosis, titer, and product quality (Bazaz et al., 2023; Leroux et al., 2021). Studies have revealed that specific combinations of miRs can result in phenotypic changes (Cursons et al., 2018; Kao et al., 2022), suggesting that the highly abundant miRs identified in CHO EVs may need to be examined in combination in order to identify any potential phenotypes.

The practical implications of this study are twofold. Differential miR and protein cargo loading suggest a use for CHO EVs as a culture diagnostic tool; indeed, the use of EV concentration as a metric for CHO culture health has already been demonstrated (Zavec et al., 2016). Both RT-PCR and our Qubit (total miR) and Bradford protein assays provide proof-of-concept for the relatively quick and simple evaluation of EV cargo. In several cases, these assays not only identify the presence of stress, but also differentiate between types of stress (i.e., ammonia or lactate vs. osmotic stress). Some EV cargo characteristics—specifically, total miR and miR-92a/let-7c levels—could even serve to differentiate between different levels of the same stress (i.e., 20 mOsm/L and 120 mOsm/L osmotic stress). Beyond their potential as a diagnostic tool, CHO EVs offer an efficient miR (or protein) delivery system that does not require disruption of the plasma membrane or complex genetic engineering. Manipulating cellular phenotype via miR is particularly appealing for CHO culture, as miRs do not compete with the translational machinery used in protein production (Muller, Katinger, & Grillari, 2008). In this vein, this study reports novel findings regarding the pro- or anti-apoptotic functions (in CHO cells) of let-7a, miR-21, and miR-92a, and also provides a framework for understanding the age/stress conditions that produce CHO EVs with high or low levels of certain miRs.

Future research should investigate the identities of the specific protein chaperones responsible for altered EV cargo loading under stress. Additionally, though miR-92a and miR-21 are highly expressed by CHO MPs from early exponential phase culture and let-7a is highly expressed by CHO MPs produced under stress (Belliveau & Papoutsakis, 2023), these miRs are by no means the only cargo capable of inducing phenotype changes in target cells. The roles of other miRs—especially those present at high levels in EVs from stress conditions—must be evaluated. At some point, it will be necessary to conduct RNA sequencing (and possibly proteomic analysis) of CHO EVs produced under lactate stress and in lag- and stationary-phase culture, as we predict the presence of novel EV miR profiles in these conditions. In sum, our research initiates the long process of linking CHO EV cargo with CHO EV function, suggesting exciting applications for CHO EVs as both diagnostic tools and culture therapeutics.

## AUTHORSHIP CONTRIBUTIONS

JB, WT and ETP designed the study and analyzed the data. JB and WT carried out the experiments. JB, WT and ETP wrote the manuscript. JB and WT should be considered joint first author.

## CONFLICTS OF INTEREST

The authors declare no conflicts of interest.

## FUNDING

This work was supported by Merck & Co., Inc. and the Advanced Mammalian Biomanufacturing Innovation Center (AMBIC). WT was additionally funded by a Department of Education GAANN Fellowship (grant number P200A210065)

## DATA AVAILABILITY

The data used to support the findings of this study are available from the corresponding author upon reasonable request.

## Notes

### Competing Interest Statement

The authors have declared no competing interest.

